# Synergistic Interactions Between Black Soldier Fly Larvae and *Thiobacillus thioparus Beijerinck 1904* for Ammonia Odor Control in Food Waste Bioconversion

**DOI:** 10.64898/2026.05.04.722119

**Authors:** Jitao Fang, Masami Shimoda

## Abstract

Black Soldier Fly larvae (BSFL, *Hermetia illucens*) are highly effective for the bioconversion of food waste. However, their rearing process often produces substantial ammonia emissions, which are malodorous and environmentally concerning. We investigated the co-cultivation of BSFL with the sulfur-oxidizing bacterium *Thiobacillus thioparus* as a strategy to mitigate ammonia release. Importantly, under conditions where ammonia emissions were significantly reduced, neither larval growth nor bacterial viability was negatively affected. Furthermore, even when the initial bacterial inoculum was reduced to 3.3*105 CFU/g-food wastes, the bacterium rapidly recovered to functional levels and effectively controlled ammonia emissions. This indicates the absence of harmful interaction or nutrient competition between BSFL and *T. thioparus*. These findings suggest an efficient method for controlling ammonia in large-scale BSFL waste treatment. By reducing the required bacterial inoculum, this approach enables scalable microbial co-culturing with environmental and production benefits.

## 1. Introduction

Organic waste management has emerged as a global environmental challenge due to the rapid increase in its generation. The accelerating growth of the world population, projected to reach 9.7 billion by 2050 (Danny Dorling 2021). Factors such as rapid urbanization, accelerated economic growth, and population expansion have placed severe pressure on waste management systems, often rendering conventional treatment methods insufficient. In particular, the treatment of household food waste has become extremely challenging with the progress of urbanization. Traditional approaches, including landfilling, incineration, and open dumping, can be effective as short-term solutions but impose substantial environmental burdens in the long term and cannot be regarded as sustainable waste management measures (Hefa Cheng, Yuanan Hu 2010; L. Giusti 2009; Daniela Porta et al.2009).

Black soldier fly *(Hermetia illucens* L., Diptera: Stratiomyidae), has attracted considerable attention as a sustainable solution to organic waste management challenges (D. Purkayastha1, S. Sarkar 2022; Qihang Zhang et al.2025). Black soldier fly larvae (BSFL) can digest a wide range of organic waste and convert it into valuable resources through their metabolic processes, enabling the sustainable recycling of food waste and livestock manure through bioconversion (Lorenzo Mazza et al. 2020; Junhua Ma et al. 2018; Trinh T. X. Nguyen et al. 2015). However, the large-scale application of BSFL in food waste treatment faces critical challenges, particularly regarding odor emissions. Ammonia is recognized as one of the most significant contributors to offensive odors and gaseous emissions during waste treatment (Alejandro Parodi et al. 2020; Shuguang Wang, Yang Zeng 2018). Beyond odor concerns, nitrogen loss through ammonia volatilization during composting reduces the overall recycling efficiency of organic waste (Dennis Beesigamukama et al. 2020; Hong Giang Hoang et al. 2022), representing both an environmental and resource conservation issue. Addressing ammonia emissions is therefore crucial for improving the sustainability and feasibility of BSFL-based food waste treatment systems.

Understanding the role of microbial communities in BSFL rearing systems may provide insights into mitigating these challenges (Jeroen De Smet 2018). Recent studies on BSFL-mediated food waste bioconversion have revealed that the dominant microbial communities in BSFL-rearing environments are highly variable rather than fixed. Distinct microbial assemblages have been reported across different studies, with compositions largely dependent on the regional source and characteristics of the food waste used. In some cases, these variations have even led to contradictory outcomes regarding system performance (Cheng-Liang Jiang et al. 2019; Moritz Gold et al. 2020). These findings suggest that BSFL possess limited intrinsic capacity to regulate or stabilize their surrounding microbiota and that their development does not rely on obligatory symbiotic microorganisms (Xin-Yu Li et al. 2019; Shu-Wei Lin, Matan Shelomi 2024). This ecological flexibility presents a unique opportunity: the microbial community within BSFL rearing environments can potentially be deliberately manipulated to achieve desired functional outcomes, such as mitigating ammonia emissions during the treatment of food waste (Lusheng Li et al. 2023).

Among microbial candidates for addressing this issue, *Thiobacillus thioparus* has drawn considerable attention due to its unique sulfur-oxidizing capability (Luc Malhautier et al. 2003). *T. thioparus* is a motile rod-shaped bacterium, with optimal growth occurring at 25–30°C and pH 6.0–8.0 conditions similar to those preferred by BSFL (Wancheng Pang et al. 2020). Owing to its ability to oxidize organic and inorganic sulfur compounds, *T. thioparus* has been widely applied in biofilters to suppress hydrogen sulfide emissions (Patricio Oyarzún et al. 2003; Heesung Kim et al. 2002). In peat-based biofilters, it has been demonstrated that *T. thioparus* can remove ammonia through chemical reactions between NH₃ and H₂SO₄ (Wenjie Gu et al. 2018; Michael J. Gibson et al. 2006). Furthermore, research reported that the addition of only 0.25% sulfur significantly enhanced the ability of *T. thioparus* to inhibit ammonia emissions (Yusheng Lu et al. 2018). In our preliminary studies, inoculation with BSFL was shown to suppress the formation of volatile sulfur compounds (VSCs) during the composting of food waste, which consequently led to an increase in sulfur concentration in the substrate (Rena Michishita et al. 2023).

This study aimed to evaluate the synergistic effects of BSFL and *T. thioparus* on ammonia reduction during food waste conversion. Specifically, we examined the effect of sulfide emission suppression by BSFL—resulting in elevated sulfide levels in the substrate—on the ammonia inhibition capacity of *T. thioparus*. The growth of both BSFL and bacteria was monitored, and ammonia emissions were measured throughout the conversion process. Our findings provide new insights into the integration of insect-based bioconversion with microbial interventions as a sustainable strategy for mitigating nitrogen loss and odor emissions in organic waste management.

## 2. Materials and methods

### 2.1. Strain

*Thiobacillus thioparus Beijerinck 1904* was from National Institute of Technology and Evaluation (NITE) Japan, and the colony was maintained at the Laboratory of Applied Entomology, The University of Tokyo. For long-term preservation, cultures were mixed with 10% glycerol and stored at −80°C (Rich Boden et al. 2019).

Experimental BSFL used in this study were originally collected in Kagoshima, Japan, in 2020, and the colony was maintained at the Laboratory of Applied Entomology, The University of Tokyo, following established protocols. Hatched larvae were reared in plastic cups (inner diameter 130 mm × height 100 mm, 860MB, Mineron Kasei Co., Osaka, Japan) and fed a diet of breadcrumbs and rice bran. Prepupae were collected at the prepupal stage and transferred to new plastic containers filled with wood chips for pupation. The containers were placed in insect-proof cages and maintained in a biotron under a 16L:8D light cycle at 32.5 ± 2.5°C and 60 ± 10% relative humidity and were provided with water (Hiroto Ohki et al. 2025).

### 2.2. Media formulations

A solution of KH₂PO₄ 1.8 g, Na₂HPO₄ 1.2 g, (NH₄)₂SO₄ 0.1 g, MgSO₄·7H₂O 0.1 g, FeCl₃·6H₂O 30 mg, MnSO₄·H₂O 30 mg, and CaCl₂·2H₂O 40 mg in 900 mL distilled water was adjusted to pH 7 with sodium bicarbonate, autoclaved at 121 °C for 20 min, and supplemented under aseptic conditions with 100 mL of sterile-filtered 10%Na₂S₂O₃ solution. To enhance solubility and minimize precipitation, the conventional S6 medium was modified by substituting some compounds with their hydrated forms. (M. Hutchinson et al. 1965). For agar medium, 15 g of agar was added to the same base solution before autoclaving (Kim D. Jones et al. 2012). The prepared liquid and solid media were stored at 4 °C. Media stored for more than one month exhibited a markedly decreased cultivation efficiency.

For research, 50 mL of the liquid medium was added to a 300 mL Erlenmeyer flask. A frozen *T. thioparus* stock (150 µL, stored at −80 °C) was thawed and inoculated into the liquid medium. The flasks were sealed with cotton plugs and incubated at 30 °C under aerated conditions with a 16 L:8 D light cycle (Rich Boden et al. 2019). The optical density (OD) of the culture was measured every 12 h to construct a growth curve.

### 2.3. Artificial food waste

In the laboratory, an artificial food waste was formulated to reflect the composition of a typical Japanese household, including five food groups: vegetables, fruits, carbohydrates, meat, and fish (Supplementary Table 1; Food Waste Suitable for Treatment Using Black Soldier Fly Larvae). Prior to introducing the larvae, freshly prepared waste was left to partially decompose for 3 days (Rena Michishita et al. 2023).

### 2.4. Experimental design

Four treatments were established in this study: (1) food waste only (T1), (2) food waste with larvae (T2), (3) food waste with *T. thioparus* suspension (T3), and (4) food waste with both larvae and *T. thioparus* suspension (T4). Each treatment contained 100 g of food waste, 30 black soldier fly larvae, and 2% (v/w) *T. thioparus* suspension, which were added when applicable. The *T. thioparus* suspension was evenly sprayed onto the food waste and thoroughly mixed. Newly hatched larvae were fed an artificial diet containing glucose, molasses, yeast, p-hydroxybenzoic acid methyl ester, propionic acid, agar, distilled water and cornmeal (Supplementary Table 2; artificial diet Suitable for Black Soldier Fly Larvae) (C.-M. Liu et al. 2025) for seven days prior to the experiment to ensure a relatively uniform body weight before introduction and a high survival rate after being added to the food waste. The prepared mixtures were incubated in plastic containers (130 mm diameter × 100 mm height, 860MB) at 30°C under a 16L:8D light cycle with ventilation for seven days. The lids of the plastic containers were perforated with 50 holes (2 mm in diameter) using a conical needle to allow aeration. Each day, the containers were removed from the incubator and placed in a well-ventilated area for 5 min to minimize mutual interference among samples before measurement. Ammonia concentrations were then measured using a GX-6000 gas detector (Riken Keiki Co., Ltd., Japan) at a height of 2 cm above each sample for 5 min. The highest value recorded during this period was defined as the ammonia concentration associated with odor intensity. On each day, five samples (0.2g each) were collected from five different locations within the container and homogenized for analysis, all samples were stored at −20 °C until further use (Rena Michishita et al. 2023; Wenjie Gu et al. 2011). In treatments containing larvae, fifteen larvae were randomly collected daily to measure the mean body weight for constructing growth curves. On the final day, all 30 larvae were collected and weighed to evaluate the effect of *T. thioparus* on BSFL growth, and the frass was also collected and sent to Shimadzu Corporation (Kawasaki, Japan) for metabolite analysis using a GCMS-8040 Triple Quadrupole Gas Chromatograph Mass Spectrometer (TQ).

### 2.5. DNA extraction

Total DNA was extracted from the samples using the ISOFECAL DNA Extraction Kit (Nippon Gene Co., Ltd., Japan) following the manufacturer’s instructions. Briefly, 0.1 g of sample was placed in a 1.5 mL microcentrifuge tube and 0.5 mL of Lysis Solution F was added. Samples were resuspended, vortexed for 1 min, and incubated at 65 °C for 1 h. After centrifugation at 12000rpm for 5 min at room temperature, 300 μL of supernatant was transferred to a new tube and mixed with 200 μL of Purification Solution, followed by the addition of 300 μL of chloroform. The mixture was vortexed briefly and centrifuged at 12000rpm for 15 min. The aqueous phase (400 μL) was carefully transferred to a new tube, avoiding the interphase, and combined with 400 μL of precipitation solution. This mixture was then centrifuged at 4 °C. The pellet was washed with 0.5 mL of Wash Solution, centrifuged at 12000rpm for 10 min at 4 °C, and subsequently treated with 0.5 mL of 70% ethanol and 2 μL Ethachinmate. After centrifugation at 12000rpm for 5 min at 4 °C, the supernatant was removed, and the pellet was air-dried and resuspended in 50 μL TE buffer (pH 8.0). All DNA samples were stored at −20 °C until further use.

### 2.6. Primer design and QPCR analysis

Degenerate primers targeting *T. thioparus* were designed based on the 16S rRNA sequence of *T. thioparus* ATCC 815816S (GenBank accession no. M79426) and other related 16S sequences. The primer sequences were: [QYF: TGA GGG GGA AAG TGG GGG AT; QYR: GTA GGC CAT TAC CCC ACC AAC] Primer specificity was confirmed using BLAST against the NBCI bacteria (taxid:2) database (V.L. Barbosa et al. 2006; Michael J. Gibson et al. 2006). Quantitative real-time PCR (qPCR) was performed on a QuantStudio 3 instrument (Thermo Fisher Scientific, USA) using THUNDERBIRD Next Probe qPCR Mix (TOYOBO, Japan). The qPCR cycling conditions followed the manufacturer’s universal protocol: an initial denaturation at 95 °C for 20 s, followed by 40 cycles of 95 °C for 1 s and 60 °C for 1 min.

### 2.7. Construction of the standard curve analysis for real-time PCR

The population of *T. thioparus* was determined using the serial dilution and spread plate method (Janeta Starosvetsky et al. 2013). For qPCR analysis, the amplification efficiency of *T. thioparus* DNA was determined by performing regression analysis between the log₁₀-transformed cell equivalents or DNA concentrations of a series of diluted DNA samples and their corresponding Cₜ values. The detection sensitivity of the *T. thioparus* qPCR assay was further estimated from the regression line of the dilution series (Nichole E. Brinkman et al. 2003; Richard A. Haugland et al. 2005).

### 2.8. Measurement of the degradation of *T. thioparus* 16S rDNA

*T. thioparus* suspension was sterilized by autoclaving at 121 °C for 20 min. Then, 70 µL of the sterilized bacterial suspension, 3 g of food waste, and one larva were placed in a 50 mL centrifuge tube and incubated at 30 °C. At 0 min, 10 min, 20 min, 40 min, 1 h, 2 h, 3 h, 4 h, and 6 h, one centrifuge tube was removed, and 30 mL of distilled water was added. After vigorous vortexing, 0.2 mL of the suspension was collected for DNA extraction. The extracted DNA was analyzed by qPCR to determine the degradation of *T. thioparus* 16S rDNA.

### 2.9. Confirmation of *T. thioparus* survival in the gut of BSFL

BSFL reared on food waste(T2) and food waste with *T. thioparus*(T4) were collected on the final day. The larvae were starved for 24 h to minimize the influence of gut contents, then washed three times with 70% ethanol and three times with sterile water. Under sterile conditions, the guts were dissected, and total DNA was extracted. The presence of *T. thioparus* was confirmed by amplifying its 16S rDNA using the primers [LQYF: GGG TGA GTA ATG CGT CGG AA; LQYR: GTT CAA AAT GCC ATT CCC AGG T] Primer specificity was confirmed using BLAST against the NBCI bacteria (taxid:2) database. PCR amplification was performed under the following conditions: 35 cycles of denaturation at 94 °C for 30 s, annealing at 55 °C for 30 s, and extension at 72 °C for 1 min. The PCR products were examined by 2% agarose gel electrophoresis to verify the expected band size (Cheng-Liang Jiang et al. 2019; Xin-Yu Li et al. 2019).

## 3. Results and discussion

### 3.1. Growth curve of *T. thioparus* in medium

As shown in Figure 1, due to initial adaptation to the new environment, *T. thioparus* exhibited a prolonged lag phase of approximately 48h. Consequently, the biosynthesis of the required inducible enzymes and cytoplasmic components also required additional time. The exponential growth phase began around 60h, during which enzyme activity was high and metabolism accelerated, resulting in the maximum growth rate of *T. thioparus*. As nutrients were depleted and metabolic by-products accumulated and degraded in the environment, the growth rate slowed by 192h. This growth pattern is consistent with the observations reported by Gu et al. For subsequent co-culture experiments, *T. thioparus* cultures aged 72-192h will be used to ensure optimal growth efficiency and viable cell numbers (Wenjie Gu et al. 2011).

**Fig. 1.**
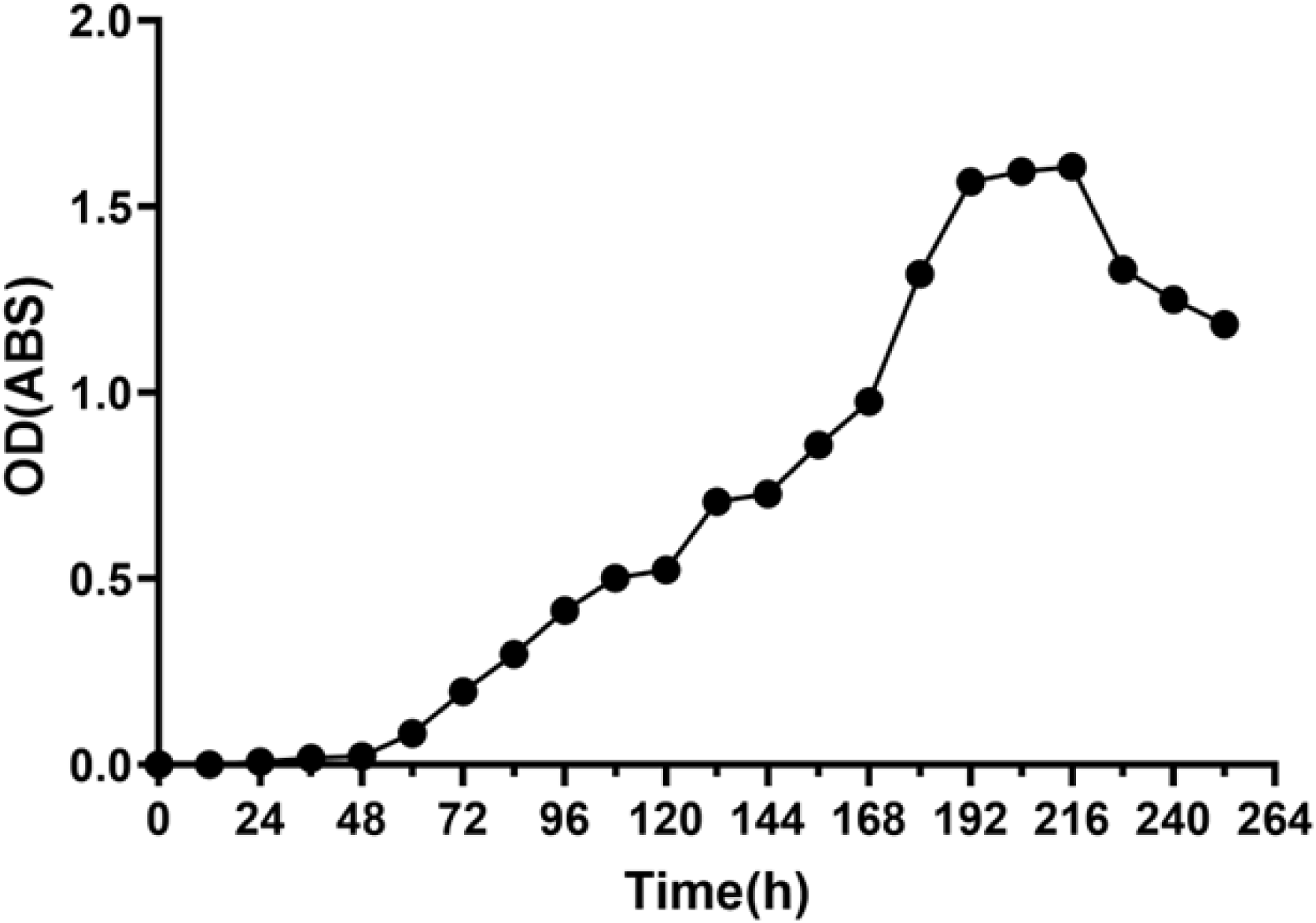
*Growth of Thiobacillus thioparus* in medium. Changes in optical density (OD) of *T. thioparus* in the culture medium over time.

### 3.2. Effect of BSFL and *T. thioparus* on ammonia-associated odor emissions from food waste

As shown in Figure 2A, the addition of BSFL in treatments T2 and T4 accelerated NH₃ emissions, whereas in treatments without BSFL (T1 and T3), ammonia release was not detected until the second day. This may be because microbial composting requires time to initiate, while the digestive enzymes (Wontae Kim et al. 2011). secreted from the salivary glands and gut of the larvae during feeding promote nitrogen mineralization, thereby increasing the concentration of ammonium (NH₄⁺) in the residual food waste (Terrence R. Green, Radu Popa 2012).From the second day onward, consistent with previous studies (Wancheng Pang et al. 2020), the T3 group, which contained only BSFL, exhibited suppression of ammonia emissions compared with T1 (food waste only) (P < 0.05). This may also be partly due to substrate reduction caused by larval feeding.

**Fig. 2.**
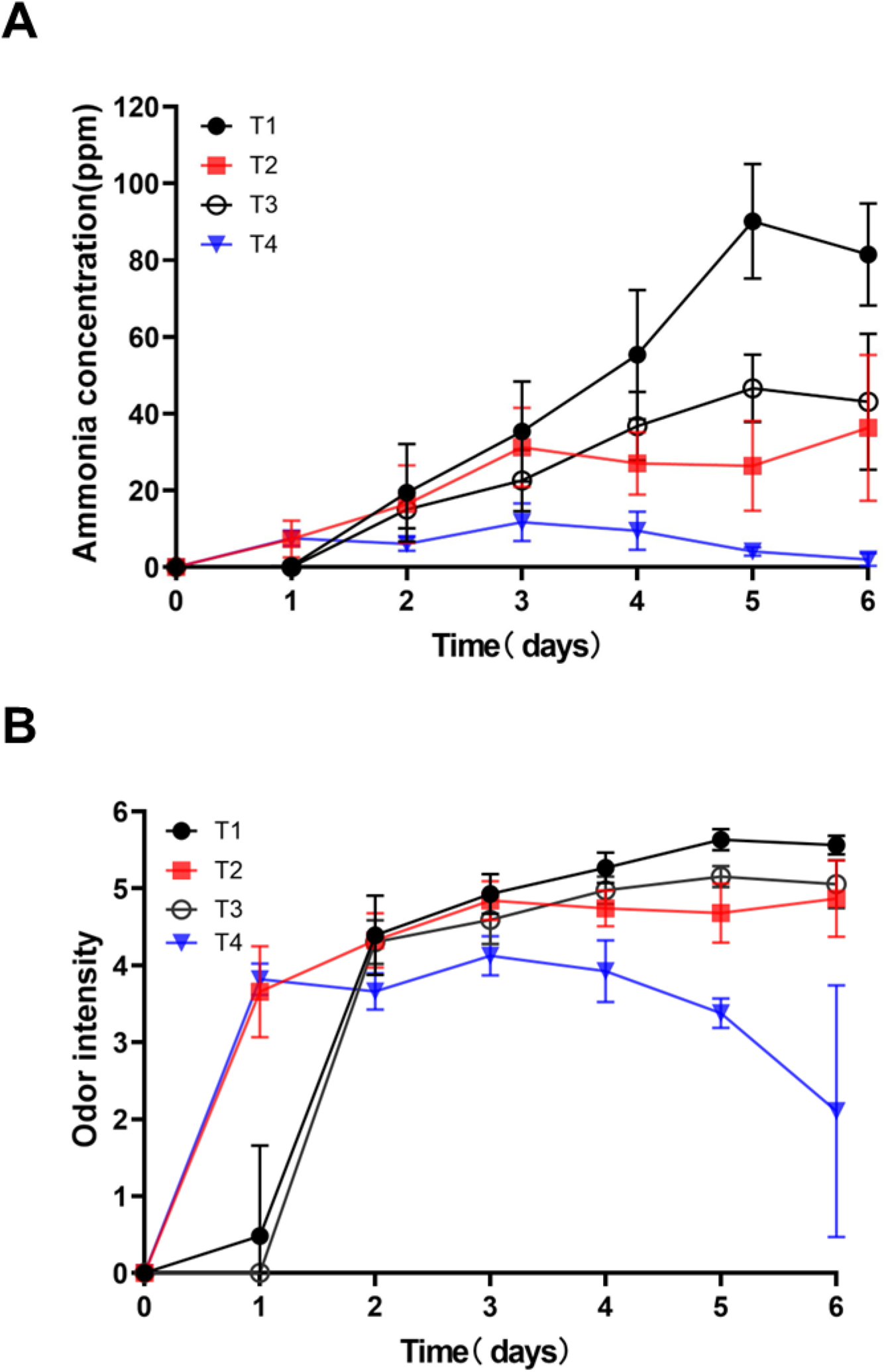
Effect of different treatments on ammonia emission from food waste substrate. (A) Temporal changes in aimnonia concentration tmder different treatments dming the experimental period. (B) Odor intensity calculated by conve1ting aimnonia concentrations at the conesponding time points for each treatment using the equation I = 1.67 log C + 2.38, where I represents odor intensity and C represents aimnonia concentration.

Regarding *T. thioparus*, we observed that in the absence of BSFL, the addition of *T. thioparus* (T3) slightly inhibited ammonia production compared with T1. However, when combined with BSFL, the T4 treatment significantly reduced the ammonia emission rate (P < 0.05). These results are consistent with the findings of Gu et al. (Wenjie Gu et al. 2018), showing that while the addition of *T. thioparus* alone can slightly reduce ammonia emission rates, the combined addition of BSFL and *T. thioparus* can effectively suppress ammonia release. This observation partially confirms our hypothesis that during the early stage of composting, proteins in meat and fish within the food waste degrade first, releasing sulfur compounds (D. Dave A.E., Ghaly 2011; Ruo He et al. 2018). The sulfur content in the meat exceeds the 0.25% sulfur addition used in previous studies. BSFL feeding suppresses the volatilization of these sulfur compounds, providing an opportunity for *T. thioparus* to oxidize organic and inorganic sulfur compounds into SO₄²⁻, which can combine with NH₄⁺ to form more stable compounds, thereby reducing ammonia emissions.

As shown in Figure 2B, ammonia concentrations were converted into odor intensity based on the Weber–Fechner law (JOANNA KOŚMIDER, BARTOSZ WYSZYŃSKI 2002) and Japanese governmental guidelines reported online (Environment Bureau of Osaka City Govt 2025) (I = 1.67 log C + 2.38, where I represents odor intensity and C represents ammonia concentration). Regardless of BSFL addition, the odor intensity far exceeded the comfort threshold of 3 for humans (Supplementary Table 3). However, co-cultivation of BSFL and *T. thioparus* not only markedly reduced ammonia concentrations but also suppressed the odor intensity caused by ammonia to below 4, making ammonia-related odor during BSFL production more acceptable.

### 3.3. Effect of *T. thioparus* on the growth of BSFL

Figure 3 shows the growth performance of BSFL co-cultured with *T. thioparus*. During the experimental period, the larval body weights in both treatments (T2 and T4) increased continuously, displaying typical sigmoidal growth curves (Fig. 3A). No significant difference (p > 0.05) was observed between the two treatments, indicating that the presence of *T. thioparus* did not significantly affect larval growth. During the growth process, half of the larvae were randomly sampled each day for body weight measurement, and their average values were used for analysis. To minimize the effects of individual size variation since — larger individuals are more likely to be found and collected in the substrate, all larvae were collected and weighed at the end of the experiment. The average individual body weight was 0.16 ± 0.06 g (n = 90) in the T2 group and 0.17 ± 0.05 g (n = 89) in the T4 group (Fig. 3B). Only one larva died in the T4 group, while all others survived, suggesting that the co-culture condition was suitable for larval development. Although the larvae in the co-culture group were slightly heavier, statistical analysis showed no significant difference (p > 0.05)

**Fig. 3.**
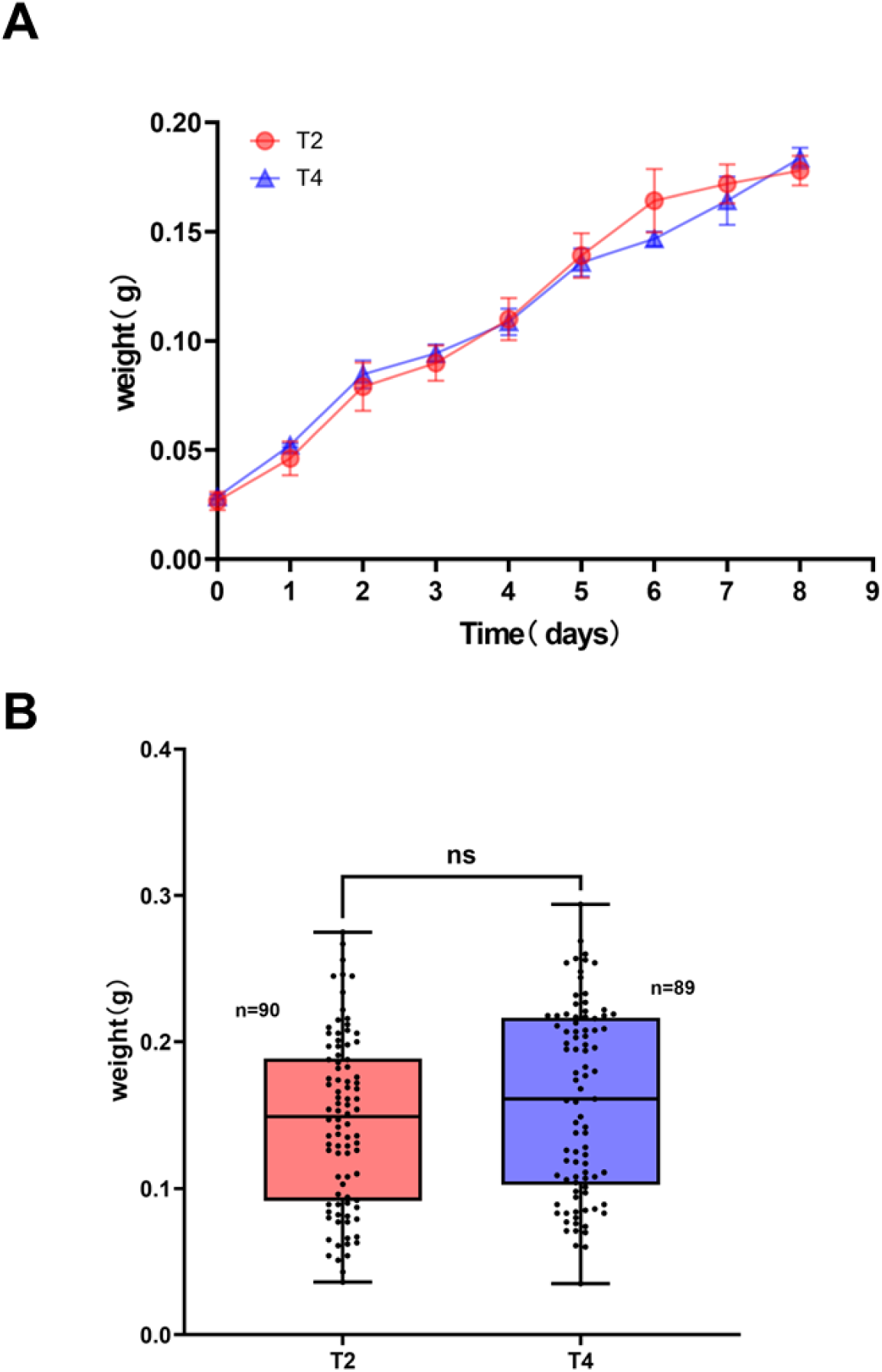
Effect of *T. thioparus* on the growth of black soldier fly larvae. (A) Daily changes in larval body weight under treatments T2 and T4 over the experimental period. For each time point, 15 larvae were randomly collected from each replicate, and the mean body weight was calculated. Data are presented as mean± SD from three independent replicates. No significant differences were detected between treatments at any time point *(p* > 0.05).(B) Body weight distribution of all larvae on the final day of the experin1ent. Each dot represents an individual larva (T2: n = 90; T4: n = 89). Box plots indicate the median, interquaitile range, and minimum-maximum values. No significant difference was observed between treatments *(p* > 0.05)

These results indicate that under the tested conditions, co-culturing with *T. thioparus* did not enhance larval growth; however, the metabolic activity of *T. thioparus* did not harm the larvae either. The similar weight gain patterns and high survival rates in both treatments suggest that nutrient availability in the food waste substrate was the main factor determining larval growth performance, and that *T. thioparus* did not compete for nutrients with the larvae. Although *T. thioparus* effectively suppressed ammonia emissions in the co-culture system, its activity did not affect larval productivity. This finding suggests that inoculating *T. thioparus* to mitigate odor emissions does not negatively impact the growth of BSFL. Consistent with the hypothesis, BSFL exhibit high environmental adaptability, and the introduction of suitable exogenous microorganisms—even those altering the rearing environment to some extent—does not adversely affect larval performance. This finding supports the potential application of BSFL–microbe co-culture in integrated and sustainable food waste bioconversion systems.

### 3.4. Quantitative PCR Detection and PCR Detection Performance Evaluation

Figure 4A shows the standard curve generated from tenfold serial dilutions of DNA extracted from food waste inoculated with *T. thioparus* (3.33 × 10⁹ CFU/mL), along with the corresponding regression equation and coefficient of determination (R²=0.9833). Based on the first standard dilution that failed to amplify in at least one replicate (10⁻⁵ dilution in the figure), the Cq cutoff value was set at 36.5. In this study, for the limiting dilution of 10⁻⁵, two out of three replicates were negative. The limit of detection (LOD) was calculated from the obtained Cq values and was determined to be <3.33 × 10^6^ CFU per gram of sample (Bojan Papić et al. 2017; Sungwoo Bae, Stefan Wuertz 2009).

**Fig. 4.**
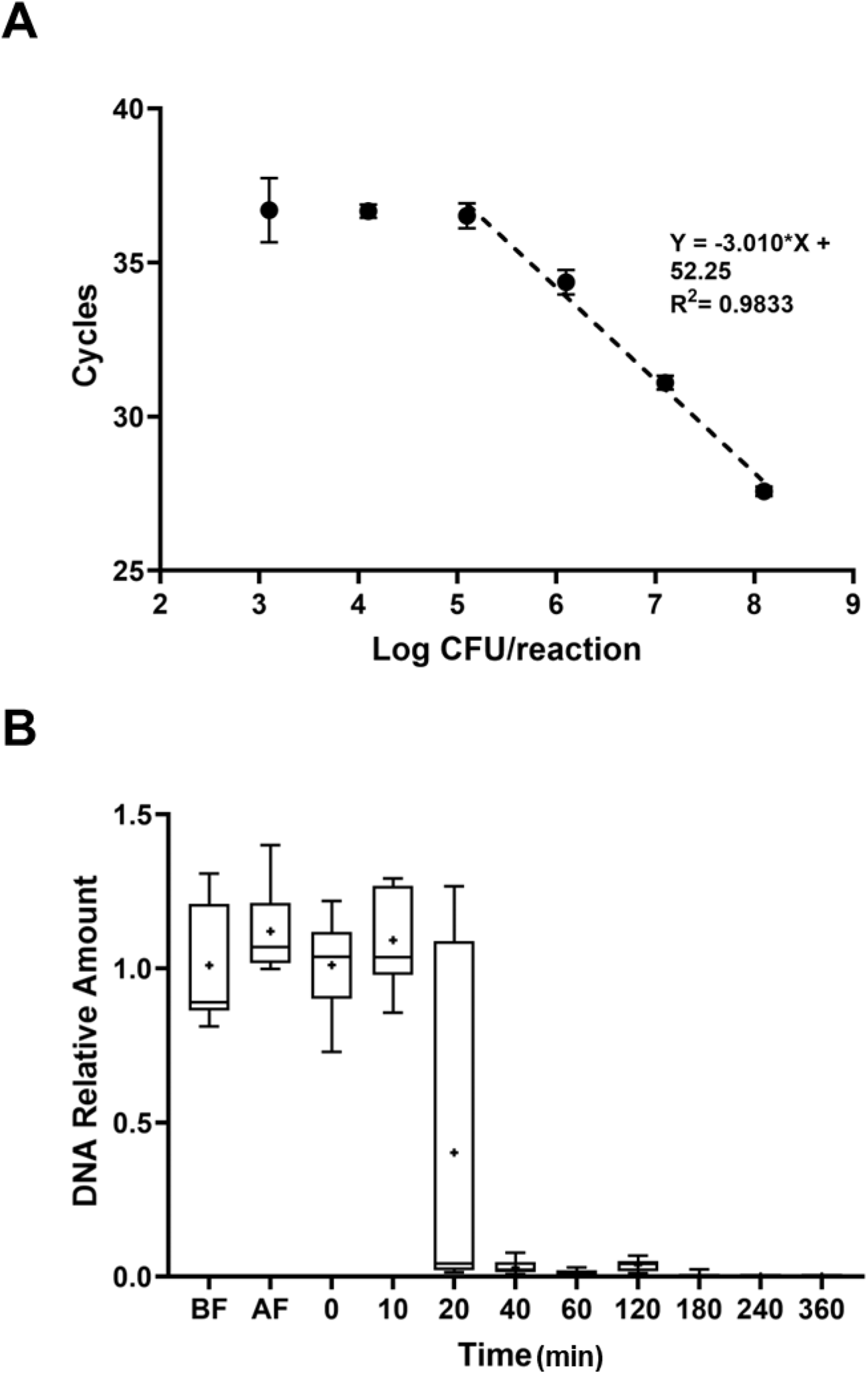
Quantification and degradation analysis of Thiobacillus thioparus 16S rDNA. (A) qPCR standard cmve showing the relationship between Ct values and log-transfonned cell numbers (log CFU). The regression equation and con-elation coefficient (R^2^) are indicated. (B) Measmement of the degradation ofT. thioparus 16S rDNA over time, quantified by qPCR. Box plots represent the distribution of relative DNA ammmts (n = 9). BF and AF represent samples collected before and after autoclaving, respectively. The “+” symbol indicates the mean value, the horizontal line inside each box represents the median, and the boxes and whiskers indicate the interquaitile range and minimum-maximum values, respectively.

**Fig. 5.**
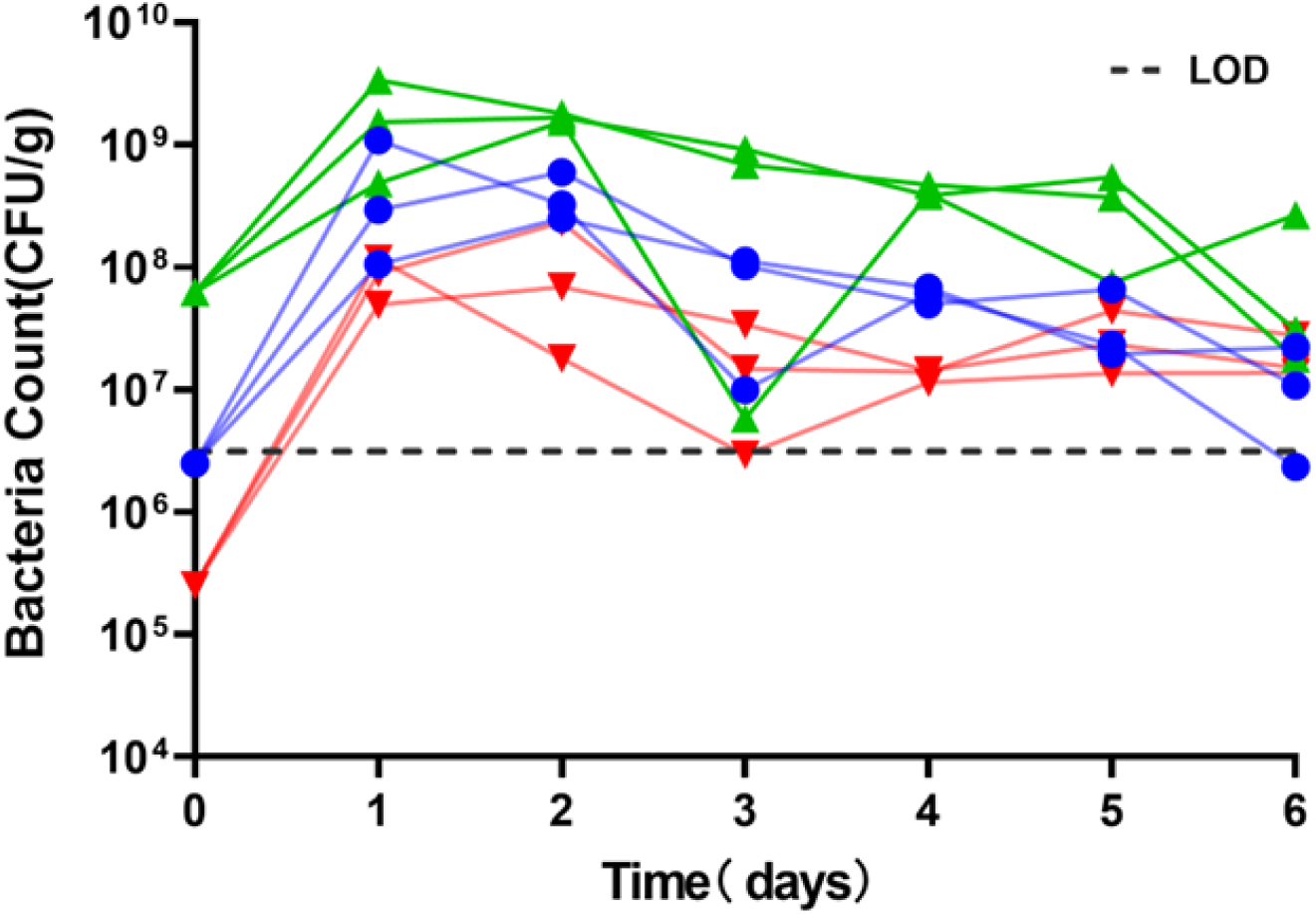
*Growth of Thiobacillus thioparus* in food waste. Temporal changes in *T. thioparus* 16S rDNA abundance in T4 over the experimental period. Each line represents an independent replicate (n = 9), with different colors indicating different initial bacterial concentrations.

To evaluate whether dead cells could interfere with the daily growth curve measurements, we investigated the degradation of *T. thioparus* 16S rDNA in a complex food waste environment following autoclaving (Geoffrey Young et al. 2007). Samples were collected before autoclaving (BF), immediately after autoclaving (AF), and at various time points within 360 min post-treatment. DNA was extracted from each sample, and the relative abundance of *T. thioparus* 16S rDNA was quantified by qPCR. As shown in Figure 4B, autoclaving did not destroy the 16S rDNA of *T. thioparus*. During the first 10 min after autoclaving, the DNA concentration remained relatively stable, comparable to the BF and AF controls. However, between 20 and 40 min, the DNA level decreased sharply from approximately 1.0 to less than 0.05 of the initial amounts. From 40 to 120 min, the DNA concentration remained negligible (relative abundance <0.1), and by 180 min it was completely degraded below the limit of detection (LOD). The rapid degradation kinetics observed here effectively eliminate concerns about DNA accumulation from dead cells, confirming that qPCR-based quantification can reliably track changes in viable bacterial populations over time. The rapid degradation may be attributed to several factors, including high microbial activity in the food waste (Nur Syahidah Zulkefli et al. 2019), the inherent instability of DNA once released from cellular protection (Tomas Lindahl 1993), and potential physical degradation in the complex food waste matrix (Rizal F. Hariadi et al. 2015; Sara Hope Sirois, Daniel H. Buckley 2019).

### 3.5. *T. thioparus* growth in food waste with black soldier fly larvae

To assess the growth and persistence of *T. thioparus*, we co-cultured *T. thioparus* with BSFL in food waste and monitored the population dynamics over seven days, with the primary goal of confirming the bacterial activity throughout one week. Although various reports indicate that DNA can persist for extended periods in vitro (Taner Çevik, Nazife Çevik 2025; Kaare M. NIELSEN et al. 2007), fortunately, the portion of *T. thioparus* 16S rDNA amplified by the qPCR primers degrades rapidly after cell death, dropping to less than 5% of the initial amount within 20–40 minutes (Fig. 4B). Since the growth curve was measured at 24-hour intervals, residual DNA from dead cells is unlikely to significantly affect daily measurements. At the start of the experiment, *T. thioparus* cultures of varying initial cell densities were inoculated into food waste. During the first two days, nutrients present in the bacterial inoculum promoted rapid exponential growth, resulting in a 2–3 log increase in cell numbers and reaching a peak on days 1–2. Furthermore, even when the initial bacterial inoculum was reduced to 3.3 × 10⁵ CFU g⁻¹ food waste, the bacterium rapidly recovered to functional levels and effectively controlled ammonia emissions, demonstrating strong resilience at low starting densities. This initial burst confirms that the introduced bacteria were active and metabolically competent. However, this peak was short-lived. After day 2, we observed a decline in viable cell numbers, likely due to nutrient depletion in the initial inoculum and subsequent decline may be due to the consumption of available substrates as BSFL gained weight, reducing the resources accessible to *T. thioparus*. This direct competition created a significant nutritional bottleneck, leading to a gradual reduction in bacterial abundance as the food waste was assimilated by BSFL.

Interestingly, while bacterial numbers generally declined steadily across most replicates, in a few instances where an unusually sharp drop occurred, a slight rebound in cell numbers was observed afterward. This “rebound” phenomenon is particularly noteworthy, as it strongly indicates that the decline in *T. thioparus* abundance was primarily due to nutrient limitation rather than inhibition by BSFL-derived antimicrobial compounds, such as previously reported antimicrobial peptides (Osama Elhag et al. 2022). This suggests that *T. thioparus* and BSFL are capable of coexisting. The results confirm that throughout the one-week period required for BSFL to process food waste, *T. thioparus* remained above the detection limit and retained high activity. In practical applications, the amount of food waste is typically much greater than that consumed by BSFL, so nutrient limitation-induced declines in bacterial abundance are unlikely to be a major concern. But for future studies, supplementation with additional materials, such as specific nutrient sources for *T. thioparus*, could be considered to further enhance ammonia suppression in the synergistic system.

### 3.6. Effect of *T. thioparus* and BSFL on Nitrogen Metabolism

The nitrogen content in frass is a critical determinant of its value as an organic fertilizer. In this study, the urea content in the frass of the T2 and T4 groups increased by 207% and 199.1% (Figure 6B), compared with the T1 control, indicating that the activity of BSFL substantially enhanced nitrogen retention in the frass. This is consistent with previous studies showing that BSFL accelerate nitrogen mineralization in food waste, with the nitrogen content in the residues nearly doubling that of the raw waste (Shwe S. Win et al. 2018). Interestingly, although the total urea content was similar between T2 and T4, the concentrations of urea precursors—ornithine and glutamic acid—were significantly higher in T4, increasing by 61.6% and 84.1% (Figure 6C, Figure 6D), respectively. This suggests that the addition of *T. thioparus* not only helps preserve bioavailable nitrogen but may also stimulate nitrogen assimilation pathways.

**Fig. 6.**
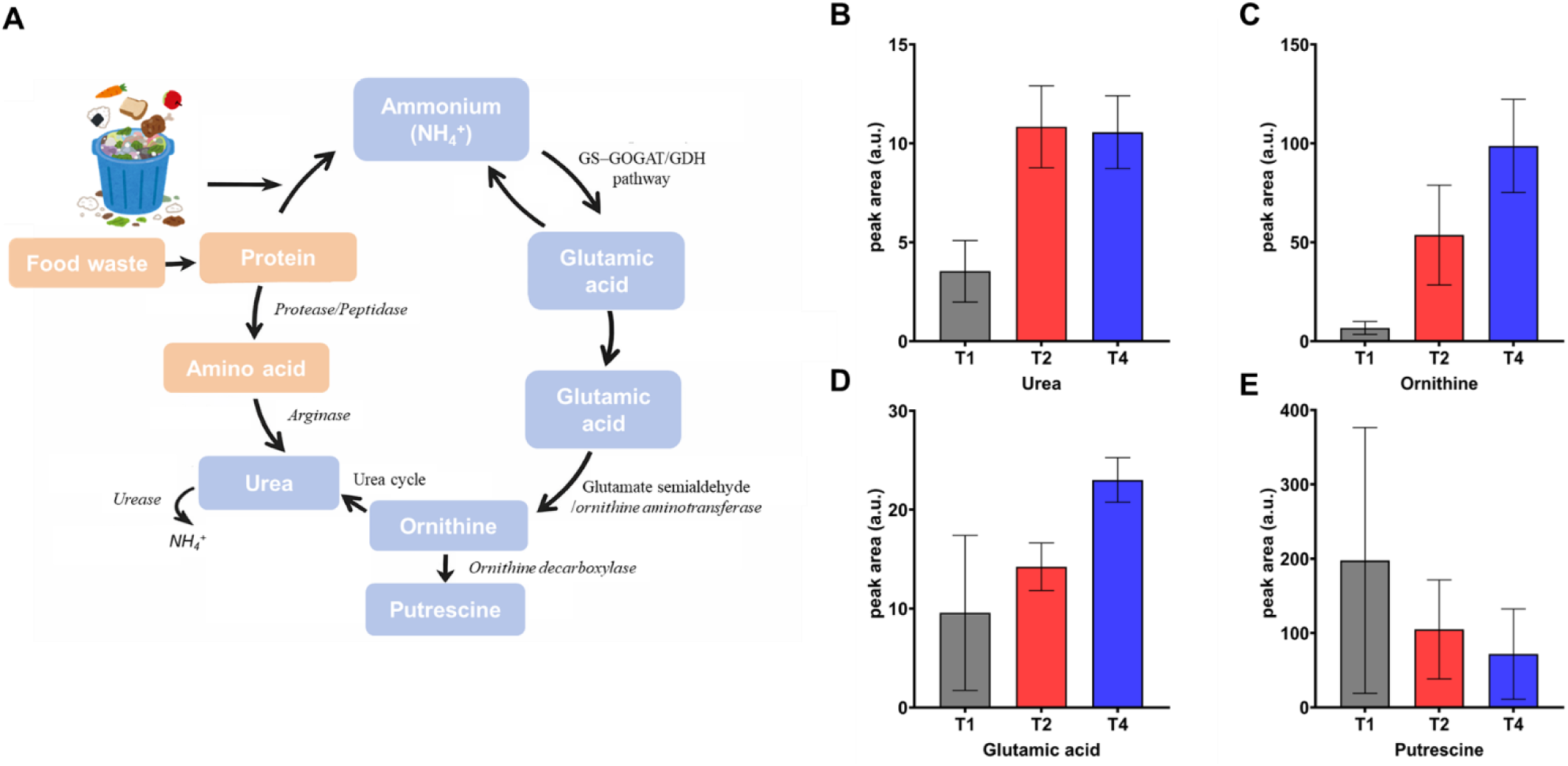
Effect of *T. thioparus* and black soldier fly larvae on icrobial nitrogen metabolism. (A) Schematic overview of microbial nitrogen metabolism pathways involved in food waste degradation, key enzymes and metabolic steps are indicate (B-E) Relative abundances of nitrogen-related metabolites measured as GC–MS peak area (a.u.) under different treatments (B) urea, (C) ornithine, (D) glutamic acid, and (E) putrescine. Data are presented as mean ± SD (n = 3).

The observed increase in precursor amino acids can be explained by coupled chemical and biological processes. BSFL activity enhances protein degradation and ammonification, leading to elevated ammonium levels in the substrate. Under such conditions, the reversible amination of α-ketoglutarate shifts toward glutamic acid formation, increasing the glutamic acid. Subsequently, sulfuric acid produced by *T. thioparus* reacts with ammonium to form ammonium sulfate, effectively reducing free ammonia volatilization while maintaining nitrogen in a biologically available form. The resulting decrease in free NH₃ concentration likely promotes assimilatory nitrogen metabolism, favoring pathways such as GS–GOGAT that incorporate ammonium into amino acids. This mechanism explains the significantly higher glutamic acid levels observed in T4 (Figure 6D) (Lin Zhu et al. 2024; ROBERT B. HELLING 1998; Jie Yuan et al. 2009). In addition, the acidification caused by H₂SO₄ accumulation can inhibit ornithine decarboxylase activity, limiting the conversion of ornithine to putrescine. Consistent with this, putrescine concentrations in T4 were markedly lower than in T2 (Figure 6E), indicating reduced ornithine catabolism and consequent ornithine accumulation (Sara Bover Cid et al. 2008).

These results indicate that synergistic interactions between BSFL and *T. thioparus* may be an effective strategy for enhancing nitrogen retention in frass. From an applied perspective, the increased urea and amino acid contents imply that frass produced under synergistic conditions may have higher fertilizer value, contributing to sustainable waste management and nutrient recycling.

## 4. Conclusions

This study demonstrates that the synergistic interaction between *Thiobacillus thioparus* and black soldier fly larvae (BSFL) effectively suppresses ammonia emissions from food waste, thereby mitigating odor during rearing without negatively affecting the growth or survival of either organism. *T. thioparus* maintained high metabolic activity throughout the seven-day bioconversion period, and even when applied at low initial inoculation levels, it rapidly proliferated to functional populations capable of controlling ammonia release, highlighting its suitability for large-scale industrial applications. BSFL activity accelerated nitrogen mineralization and significantly increased urea accumulation in the frass, while *T. thioparus* stabilized NH₃ through acid-driven conversion to NH₄⁺, further enhancing nitrogen retention. In addition, the increased ammonium availability promoted glutamate synthesis, and the lowered environmental pH inhibited ornithine decarboxylation, resulting in greater preservation and accumulation of amino acids in the frass, which contributes to improving its nutritional value when applied as an organic fertilizer.

Overall, this cooperative system reduces odor emissions, improves the fertilizer quality of the resulting frass, and maintains larval productivity. These findings provide strong evidence for integrating *T. thioparus* into BSFL-based food waste bioconversion systems and offer a sustainable strategy to optimize nutrient recycling, minimize environmental impacts, and advance the industrial application of insect-mediated waste management.

